# Optimization of the Illumina COVIDSeq^™^ protocol for decentralized, cost-effective genomic surveillance

**DOI:** 10.1101/2022.11.07.515545

**Authors:** Rob E. Carpenter, Vaibhav K. Tamrakar, Sadia Almas, Chase Rowan, Rahul Sharma

## Abstract

A decentralized surveillance system to identify local outbreaks and monitor SARS-CoV-2 Variants of Concern is one of the primary strategies for the pandemic’s containment. Although next-generation sequencing (NGS) is a gold standard for genomic surveillance and variant discovery, the technology is still cost-prohibitive for decentralized sequencing, particularly in small independent labs with limited resources. We have optimized the Illumina COVID-seq protocol to reduce cost without compromising accuracy. 90% of genomic coverage was achieved for 142/153 samples analyzed in this study. The lineage was correctly assigned to all samples (152/153) except for one. This modified protocol can help laboratories with constrained resources contribute to decentralized SARS-CoV-2 surveillance in the post-vaccination era.

---

The severe acute respiratory syndrome coronavirus (SARS-CoV-2) pandemic has claimed millions of lives globally, highlighting the need for systemic de-centralized scientific capacities. Although currently, there is a downward progression of global COVID-19 cases (Murray et al., 2022), community-level surveillance by de-centralized genomic sequencing remains vital for monitoring the transmission dynamics of the pandemic and SARS-CoV-2 mutagenic capacities. Keeping SARS-CoV-2 in the foreground, new lineages will likely emerge, and monitoring evolving variants is epidemiologically critical (Aleem et al., 2022; Anderson et al., 2021). While next-generation sequencing (NGS) has emerged as the gold standard technology for genomic surveillance and variant discovery (Berno et al., 2022), the technology remains cost prohibitive for de-centralized sequencing, particularly in small independent and resource-limited laboratories (Umunnakwe et al., 2022). Illumina COVIDSeq™ is one of the most utilized methods for COVID surveillance, but the application has been limited mainly to centralized surveillance programs.

As part of a surveillance program sponsored by the Centers for Disease Control and Prevention (CDC), 95 clinical samples from Advanta Genetics were sent to Fulgent Genetics for sequencing. Our samples were sequenced at an average depth of 20,071.41x using Illumina COVIDSeq assay on NovaSeq 6000 instrument (Cost ~ 1 million USD), which requires pooling of 1000s of samples. The standard protocol doesn’t require the normalization of libraries; thus, sequencing depth among 95 samples ranges from 286x to 287,4419× (Supplemental Table). This over-sequencing approach is acceptable in high throughput laboratories, where samples are sequenced at higher depths to achieve sufficient coverage for each sample. However, using low throughput instruments, the standard protocol is cost-prohibitive for de-centralized sequencing facilities.

Although Illumina has introduced a COVIDSeq^™^ 96 sample kit (Cat#20049393; cost/sample: $40.00), the protocol directs preparing a 50μl library from each sample and pooling only 5μl for sequencing. The standard protocol doesn’t recommend quantification and normalization of libraries before pooling, and 90% of the library volume is not used for pooling and sequencing.

Accordingly, this study describes the optimization of the Illumina COVIDSeq^™^ Research Use Only (RUO) assay protocol for the decentralized implementation of SARS-CoV-2 genomic surveillance. The protocol is modified to reduce cost and enhance efficiency on low throughput sequencing instruments like the Illumina MiniSeq^®^ or MiSeq^®^. We optimized the protocol by using 50% reagent volume at each step during library preparation, resulting in a 25μL library from each sample, cutting the library preparation cost by half ($20/sample). Next, the individual library was quantified using a Qubit™ Flex Fluorometer (Invitrogen, Inc.), and representative libraries (N=12) prepared from the reference strains were also analyzed on a fragment analyzer. The average library size (347.66±22.82 bp) and its concentration in (ng/μL) were used to calculate the molarity of the individual library. Individual libraries were also quantified using the KAPA Library Quantification Kits (Roche Cat # 07960140001). Individual libraries were diluted to achieve a normalized concentration of 10nM, and the normalized libraries were pooled in equimolar concentration instead of equal volume as recommended by the Illumina COVIDSeq™ kit. These additional normalization steps allowed us to achieve uniform coverage of all the libraries in the pool and efficiently use a low-throughput sequencing instrument. The final library pool was again quantified using a Qubit™ Flex Fluorometer (Invitrogen, Inc.) and a PCR-based library quantification kit (Scienetix, USA). The final library pool was denatured and diluted to a 2 pM loading concentration. Dual indexed paired-end sequencing with 75bp read length was carried out using the high output flow cell (list price: $1102.00) on the Illumina MiniSeq^®^ instrument. This approach allowed us to reduce the cost of the COVIDSeq^™^ reagent (library preparation + sequencing) to $56.00 per sample with a batch size of only ~30 samples. Pre-pooling normalization allowed us to achieve uniform coverage (median depth 595×) across the samples in the pool and higher efficiency. We could sequence ~30 clinical samples (diagnostic PCR Ct<30), positive control (Wuhan-Hu-1), and no template control (NTC) with each batch of libraries on a MiniSeq high-output flow cell.

The lowest sequencing depth of >200× and 90% genome coverage were found adequate for accurate variant detection while validating this assay for clinical testing according to the CAP (College of American Pathologists) guidelines for NGS-based Laboratory Developed Test (LDT) (Carpenter et al., 2022). Importantly, we sequenced 153 samples using this modified approach, and 152/153 (99.34%) samples (PCR Ct<30) resulted in the correct variant using DRAGEN COVID Lineage (v3.5.4), and 142/153 (92.81%) samples achieved 90% genome coverage with an average depth of >200x (Supplemental Table). Only 11/154 (7%) samples are below 200x coverage; out of 11 samples < 200x coverage (Table 1), the DRAGEN COVID Lineage analysis pipeline failed in assigning the lineage of only one sample with the lowest depth. We also resequenced six samples already sequenced by another reference laboratory (Fulgent Genetics, Inc.) at extremely high coverage (>30,000X) and compared the variant identities from this modified sequencing protocol. The results confirmed that all six samples were identified to carry identical variants by both laboratories, implicating 100% accuracy in the inter-laboratory testing of this modified approach. Interestingly, 3 of the six split samples were sequenced at >50,000× coverage by Fulgent Genetics, whereas we were able to sequence the same samples at only 200X coverage with identical variant detection. This illustrates higher sequencing efficiency with pre-pooling quantification without compromising the test accuracy. Such higher efficiency is essential for the cost-effective application of this test in limited-resourced and de-centralized laboratory settings and for reference laboratories that do not have access to high throughput instruments such as Illumina NextSeq^®^ or NovSeq^®^ instruments. (Thomas et al., 2021)

**Table 1:** Comparative sequencing using the modified COVIDSeq^™^ protocol

| Sequencing Parameter | Fulgent Genetics<br>(n=95*) | Advanta Genetics<br>(n=153*) |
| --- | --- | --- |
| Mean Coverage | 30257.05 | 1132.30 |
| Median Coverage | 32148.50 | 595.00 |
| Standard Deviation (SD) | 20071.41 | 1232.19 |
| Highest Coverage (X times) | 2874419.50 | 6913.00 |
| Lowest Coverage (X times) | 268.00 | 56.00 |
| Accurately Variant Calls | 100% (95/95) | 99.34 (152/153) |

\* 6 samples were split and sequenced by both approaches.

This modified protocol can potentially empower small resource-limited laboratories to contribute to local genomic surveillance. We have adopted this modified protocol for sequencing 153 genomes from East Texas, USA, and compared the results with PCR-based variant detection (Carpenter et al., 2022). High accuracy and reproducibility of this approach have been demonstrated in validating the COVIDSeq™ assay for clinical application according to Clinical Laboratory Improvement Amendments and College of American Pathologists guidelines. This cost-effective approach can be widely adopted for low-throughput sequencing and monitoring of emerging variants of SARS-CoV-2 and further support de-centralized genomic surveillance.

## Author Contributions

Conceptualization RC and RS; Data curation SA, CR and RS; Formal analysis; RS and SA Validation RS and SA; Roles/Writing: original draft RS, SA, and VT; Writing: review & editing RC.

## Statement of Informed Consent

No conflicts, informed consent, or human or animal rights apply to this work.

## Declaration of Competing Interest

RS and RC declare that they have a financial interest in Scienetix, Inc. None of the other authors declare that they have no known competing financial interests or personal relationships that could have appeared to influence the work reported in this paper.

## Notes

### Competing Interest Statement

The authors have declared no competing interest.

