## Supplemental Table for "Optimization of the Illumina COVIDSeq^™^ protocol for decentralized, cost-effective genomic surveillance"

**Supplemental Table: SARS CoV-2 variant detection using the Next Generation Sequencing at both the Laboratory (Fulgent Genetics and Advanta Genetics)**

| **Sr. No** | **Sequenced At** | **Seq Sample ID** | **Median Coverage** | **Pango  Lineage** | **WHO label** |
| --- | --- | --- | --- | --- | --- |
| 1 | Fulgent | FT-SA99813 | 69088.9 | B.1.617.2 | Delta |
| 2 | Fulgent | FT-SA99822 | 341.6 | AY.3 | Delta (B.1.617.2-like) |
| 3 | Fulgent | FT-SA99791 | 77387.8 | AY.3 | Delta (B.1.617.2-like) |
| 4 | Fulgent | FT-SA99778 | 37341.1 | B.1.617.2 | Delta |
| 5 | Fulgent | FT-SA99782 | 54823.9 | AY.3 | Delta (B.1.617.2-like) |
| 6 | Fulgent | FT-SA99829 | 2860.5 | AY.3 | Delta (B.1.617.2-like) |
| 7 | Fulgent | FT-SA99824 | 2849.4 | AY.3 | Delta (B.1.617.2-like) |
| 8 | Fulgent | FT-SA99834 | 13183.7 | AY.3 | Delta (B.1.617.2-like) |
| 9 | Fulgent | FT-SA99827 | 27506.0 | AY.3 | Delta (B.1.617.2-like) |
| 10 | Fulgent | FT-SA99825 | 24351.7 | AY.3 | Delta (B.1.617.2-like) |
| 11 | Fulgent | FT-SA99833 | 317.0 | AY.3 | Delta (B.1.617.2-like) |
| 12 | Fulgent | FT-SA99821 | 54609.4 | AY.3 | Delta (B.1.617.2-like) |
| 13 | Fulgent | FT-SA99823 | 54978.4 | AY.3 | Delta (B.1.617.2-like) |
| 14 | Fulgent | FT-SA99826 | 15381.2 | AY.3 | Delta (B.1.617.2-like) |
| 15 | Fulgent | FT-SA99820 | 22329.0 | AY.3 | Delta (B.1.617.2-like) |
| 16 | Fulgent | FT-SA99828 | 47290.6 | B.1.617.2 | Delta |
| 17 | Fulgent | FT-SA99854 | 32397.9 | B.1.617.2 | Delta |
| 18 | Fulgent | FT-SA99847 | 44269.6 | AY.3 | Delta (B.1.617.2-like) |
| 19 | Fulgent | FT-SA99838 | 44663.3 | AY.3 | Delta (B.1.617.2-like) |
| 20 | Fulgent | FT-SA99855 | 32148.5 | AY.3 | Delta (B.1.617.2-like) |
| 21 | Fulgent | FT-SA99832 | 22923.6 | AY.3 | Delta (B.1.617.2-like) |
| 22 | Fulgent | FT-SA99862 | 29984.5 | B.1.617.2 | Delta |
| 23 | Fulgent | FT-SA99837 | 36065.8 | B.1.617.2 | Delta |
| 24 | Fulgent | FT-SA99844 | 44949.1 | AY.3 | Delta (B.1.617.2-like) |
| 25 | Fulgent | FT-SA99857 | 11912.3 | B.1.617.2 | Delta |
| 26 | Fulgent | FT-SA99772 | 32888.9 | B.1.617.2 | Delta |
| 27 | Fulgent | FT-SA99861 | 17775.7 | B.1.617.2 | Delta |
| 28 | Fulgent | FT-SA99774 | 35647.4 | B.1.617.2 | Delta |
| 29 | Fulgent | FT-SA99864 | 30891.2 | AY.3 | Delta (B.1.617.2-like) |
| 30 | Fulgent | FT-SA99853 | 34400.0 | B.1.617.2 | Delta |
| 31 | Fulgent | FT-SA99831 | 66473.2 | B.1.617.2 | Delta |
| 32 | Fulgent | FT-SA99836 | 56179.3 | B.1.617.2 | Delta |
| 33 | Fulgent | FT-SA99845 | 45936.6 | B.1.617.2 | Delta |
| 34 | Fulgent | FT-SA99848 | 20892.1 | B.1.617.2 | Delta |
| 35 | Fulgent | FT-SA99830 | 57742.9 | B.1.617.2 | Delta |
| 36 | Fulgent | FT-SA99839 | 55156.1 | B.1.617.2 | Delta |
| 37 | Fulgent | FT-SA99843 | 6701.8 | B.1.617.2 | Delta |
| 38 | Fulgent | FT-SA99851 | 31771.5 | AY.3 | Delta (B.1.617.2-like) |
| 39 | Fulgent | FT-SA99846 | 44996.0 | AY.3 | Delta (B.1.617.2-like) |
| 40 | Fulgent | FT-SA99835 | 20016.8 | AY.3 | Delta (B.1.617.2-like) |
| 41 | Fulgent | FT-SA99841 | 32880.5 | AY.3 | Delta (B.1.617.2-like) |
| 42 | Fulgent | FT-SA99842 | 27810.2 | AY.3 | Delta (B.1.617.2-like) |
| 43 | Fulgent | FT-SA99849 | 37296.9 | AY.3 | Delta (B.1.617.2-like) |
| 44 | Fulgent | FT-SA99858 | 28430.1 | AY.3 | Delta (B.1.617.2-like) |
| 45 | Fulgent | FT-SA99840 | 34747.9 | AY.3 | Delta (B.1.617.2-like) |
| 46 | Fulgent | FT-SA99850 | 5569.7 | AY.3 | Delta (B.1.617.2-like) |
| 47 | Fulgent | FT-SA99865 | 31304.7 | AY.3 | Delta (B.1.617.2-like) |
| 48 | Fulgent | FT-SA99859 | 20662.4 | AY.3 | Delta (B.1.617.2-like) |
| 49 | Fulgent | FT-SA99803 | 2394.7 | AY.3 | Delta (B.1.617.2-like) |
| 50 | Fulgent | FT-SA99793 | 36682.4 | AY.3 | Delta (B.1.617.2-like) |
| 51 | Fulgent | FT-SA99795 | 49623.0 | AY.3 | Delta (B.1.617.2-like) |
| 52 | Fulgent | FT-SA99785 | 19843.7 | AY.3 | Delta (B.1.617.2-like) |
| 53 | Fulgent | FT-SA99790 | 4566.0 | AY.3 | Delta (B.1.617.2-like) |
| 54 | Fulgent | FT-SA99798 | 61671.5 | B.1.617.2 | Delta |
| 55 | Fulgent | FT-SA99804 | 40157.8 | B.1.617.2 | Delta |
| 56 | Fulgent | FT-SA99796 | 67119.2 | B.1.617.2 | Delta |
| 57 | Fulgent | FT-SA99802 | 35788.0 | AY.3 | Delta (B.1.617.2-like) |
| 58 | Fulgent | FT-SA99792 | 2425.0 | B.1.617.2 | Delta |
| 59 | Fulgent | FT-SA99801 | 9208.1 | B.1.617.2 | Delta |
| 60 | Fulgent | FT-SA99777 | 29240.8 | AY.3 | Delta (B.1.617.2-like) |
| 61 | Fulgent | FT-SA99784 | 1029.7 | AY.3 | Delta (B.1.617.2-like) |
| 62 | Fulgent | FT-SA99771 | 35543.9 | AY.3 | Delta (B.1.617.2-like) |
| 63 | Fulgent | FT-SA99799 | 40148.8 | AY.3 | Delta (B.1.617.2-like) |
| 64 | Fulgent | FT-SA99797 | 10491.7 | AY.3 | Delta (B.1.617.2-like) |
| 65 | Fulgent | FT-SA99789 | 57325.7 | AY.3 | Delta (B.1.617.2-like) |
| 66 | Fulgent | FT-SA99783 | 33664.1 | AY.3 | Delta (B.1.617.2-like) |
| 67 | Fulgent | FT-SA99775 | 2096.2 | AY.3 | Delta (B.1.617.2-like) |
| 68 | Fulgent | FT-SA99779 | 35032.8 | AY.3 | Delta (B.1.617.2-like) |
| 69 | Fulgent | FT-SA99794 | 35808.2 | B.1.617.2 | Delta |
| 70 | Fulgent | FT-SA99787 | 41262.8 | B.1.617.2 | Delta |
| 71 | Fulgent | FT-SA99809 | 29924.4 | B.1.617.2 | Delta |
| 72 | Fulgent | FT-SA99806 | 71332.2 | B.1.617.2 | Delta |
| 73 | Fulgent | FT-SA99773 | 33387.6 | B.1.617.2 | Delta |
| 74 | Fulgent | FT-SA99819 | 6792.1 | B.1.617.2 | Delta |
| 75 | Fulgent | FT-SA99811 | 48201.1 | B.1.617.2 | Delta |
| 76 | Fulgent | FT-SA99812 | 47985.0 | B.1.617.2 | Delta |
| 77 | Fulgent | FT-SA99815 | 3672.1 | AY.3 | Delta (B.1.617.2-like) |
| 78 | Fulgent | FT-SA99814 | 70771.4 | AY.3 | Delta (B.1.617.2-like) |
| 79 | Fulgent | FT-SA99817 | 26266.6 | AY.3 | Delta (B.1.617.2-like) |
| 80 | Fulgent | FT-SA99818 | 30345.7 | AY.3 | Delta (B.1.617.2-like) |
| 81 | Fulgent | FT-SA99816 | 36172.8 | AY.3 | Delta (B.1.617.2-like) |
| 82 | Fulgent | FT-SA99863 | 358.1 | B.1.617.2 | Delta |
| 83 | Fulgent | FT-SA99860 | 2015.3 | B.1.617.2 | Delta |
| 84 | Fulgent | FT-SA99856 | 268.0 | B.1.617.2 | Delta |
| 85 | Fulgent | FT-SA99852 | 596.6 | AY.3 | Delta (B.1.617.2-like) |
| 86 | Fulgent | FT-SA99808 | 32246.4 | B.1.617.2 | Delta |
| 87 | Fulgent | FT-SA99807 | 57736.7 | B.1.617.2 | Delta |
| 88 | Fulgent | FT-SA99805 | 43664.1 | B.1.617.2 | Delta |
| 89 | Fulgent | FT-SA99780 | 6849.1 | B.1.617.2 | Delta |
| 90 | Fulgent | FT-SA99800 | 9087.8 | B.1.617.2 | Delta |
| 91 | Fulgent | FT-SA99781 | 1288.2 | B.1.617.2 | Delta |
| 92 | Fulgent | FT-SA99770 | 22127.2 | B.1.617.2 | Delta |
| 93 | Fulgent | FT-SA99788 | 49129.7 | AY.3 | Delta (B.1.617.2-like) |
| 94 | Fulgent | FT-SA99786 | 1809.3 | AY.3 | Delta (B.1.617.2-like) |
| 95 | Fulgent | FT-SA99776 | 11141.2 | AY.3 | Delta (B.1.617.2-like) |
| 96 | Advanta | ACSQ1-16 | 3739 | B.1 | Non- VOC |
| 97 | Advanta | ACSQ1-11 | 2609 | B.1.2 | Non- VOC |
| 98 | Advanta | ACSQ1-18 | 2494 | B.1.564 | Non- VOC |
| 99 | Advanta | ACSQ1-21 | 2393 | B.1.617.2 | Delta |
| 100 | Advanta | ACSQ2-1 | 2289 | AY.25 | Delta (B.1.617.2-like) |
| 101 | Advanta | ACSQ1-19 | 2044 | B.1.617.2 | Delta |
| 102 | Advanta | ACSQ1-9 | 1905 | B.1.574 | Non- VOC |
| 103 | Advanta | ACSQ1-20 | 1901 | AY.3 | Delta |
| 104 | Advanta | ACSQ1-15 | 1607 | B.1.602 | Non- VOC |
| 105 | Advanta | ACSQ1-17 | 1577 | B.1 | Non- VOC |
| 106 | Advanta | ACSQ1-13 | 1472 | B.1.234 | Non- VOC |
| 107 | Advanta | ACSQ4-7 | 1438 | B.1.1.529 | Omicron (B.1.1.529-like) |
| 108 | Advanta | ACSQ4-9 | 1363 | B.1.1.529 | Omicron (B.1.1.529-like) |
| 109 | Advanta | ACSQ1-8 | 1352 | B.1.243 | Non- VOC |
| 110 | Advanta | ACSQ4-15 | 1308 | B.1.1.529 | Omicron (B.1.1.529-like) |
| 111 | Advanta | ACSQ4-19 | 1296 | B.1.1.529 | Omicron (B.1.1.529-like) |
| 112 | Advanta | ACSQ4-1 | 1286 | B.1.1.529 | Omicron (B.1.1.529-like) |
| 113 | Advanta | ACSQ4-14 | 1273 | B.1.1.529 (probable) | Probable Omicron (B.1.1.529-like) |
| 114 | Advanta | ACSQ4-23 | 1265 | B.1.1.529 | Omicron (B.1.1.529-like) |
| 115 | Advanta | ACSQ4-18 | 1244 | B.1.1.529 | Omicron (B.1.1.529-like) |
| 116 | Advanta | ACSQ4-5 | 1227 | B.1.1.529 (probable) | Probable Omicron (B.1.1.529-like) |
| 117 | Advanta | ACSQ4-13 | 1209 | B.1.1.529 | Omicron (B.1.1.529-like) |
| 118 | Advanta | ACSQ4-11 | 1201 | B.1.1.529 | Omicron (B.1.1.529-like) |
| 119 | Advanta | ACSQ4-26 | 1166 | B.1.1.529 | Omicron (B.1.1.529-like) |
| 120 | Advanta | ACSQ4-22 | 1143 | B.1.1.529 | Omicron (B.1.1.529-like) |
| 121 | Advanta | ACSQ4-3 | 1131 | B.1.1.529 | Omicron (B.1.1.529-like) |
| 122 | Advanta | ACSQ4-6 | 1130 | B.1.1.529 | Omicron (B.1.1.529-like) |
| 123 | Advanta | ACSQ1-14 | 1111 | B.1.126 | Non- VOC |
| 124 | Advanta | ACSQ4-25 | 1105 | B.1.1.529 | Omicron (B.1.1.529-like) |
| 125 | Advanta | ACSQ4-24 | 1088 | B.1.1.529 | Omicron (B.1.1.529-like) |
| 126 | Advanta | ACSQ4-16 | 1059 | AY.103 | Delta (B.1.617.2-like) |
| 127 | Advanta | ACSQ4-12 | 999 | AY.103 | Delta (B.1.617.2-like) |
| 128 | Advanta | ACSQ4-8 | 989 | AY.103 | Delta (B.1.617.2-like) |
| 129 | Advanta | ACSQ4-10 | 900 | B.1.1.529 | Omicron (B.1.1.529-like) |
| 130 | Advanta | ACSQ4-20 | 867 | AY.3 | Delta (B.1.617.2-like) |
| 131 | Advanta | ACSQ4-2 | 867 | B.1.1.529 | Omicron (B.1.1.529-like) |
| 132 | Advanta | ACSQ2-9 | 797 | AY.3 | Delta (B.1.617.2-like) |
| 133 | Advanta | ACSQ5-7 | 794 | BA.1 | Omicron (BA.1-like) |
| 134 | Advanta | ACSQ2-22 | 786 | AY.39.1 | Delta (B.1.617.2-like) |
| 135 | Advanta | ACSQ2-3 | 773 | AY.25 | Delta (B.1.617.2-like) |
| 136 | Advanta | ACSQ2-20 | 757 | AY.39.1 | Delta (B.1.617.2-like) |
| 137 | Advanta | ACSQ5-29 | 745 | BA.1 | Omicron (BA.1-like) |
| 138 | Advanta | ACSQ4-4 | 744 | AY.103 | Delta (B.1.617.2-like) |
| 139 | Advanta | ACSQ5-30 | 736 | BA.1 | Omicron (BA.1-like) |
| 140 | Advanta | ACSQ2-10 | 731 | AY.3 | Delta (B.1.617.2-like) |
| 141 | Advanta | ACSQ2-21 | 706 | AY.39.1 | Delta (B.1.617.2-like) |
| 142 | Advanta | ACSQ5-24 | 690 | BA.1 | Omicron (BA.1-like) |
| 143 | Advanta | ACSQ2-2 | 684 | AY.25 | Delta (B.1.617.2-like) |
| 144 | Advanta | ACSQ5-23 | 663 | BA.1 | Omicron (BA.1-like) |
| 145 | Advanta | ACSQ5-5 | 641 | BA.1 | Omicron (BA.1-like) |
| 146 | Advanta | ACSQ5-22 | 635 | BA.1 | Omicron (BA.1-like) |
| 147 | Advanta | ACSQ2-17 | 633 | AY.103 | Delta (B.1.617.2-like) |
| 148 | Advanta | ACSQ5-13 | 628 | BA.1 | Omicron (BA.1-like) |
| 149 | Advanta | ACSQ2-19 | 595 | AY.103 | Delta (B.1.617.2-like) |
| 150 | Advanta | ACSQ5-26 | 586 | BA.1 | Omicron (BA.1-like) |
| 151 | Advanta | ACSQ2-18 | 583 | AY.103 | Delta (B.1.617.2-like) |
| 152 | Advanta | ACSQ5-14 | 562 | BA.1 | Omicron (BA.1-like) |
| 153 | Advanta | ACSQ2-16 | 560 | AY.3 | Delta (B.1.617.2-like) |
| 154 | Advanta | ACSQ5-6 | 550 | BA.1 | Omicron (BA.1-like) |
| 155 | Advanta | ACSQ2-14 | 547 | AY.3 | Delta (B.1.617.2-like) |
| 156 | Advanta | ACSQ5-1 | 536 | BA.1 | Omicron (BA.1-like) |
| 157 | Advanta | ACSQ5-27 | 533 | BA.1 | Omicron (BA.1-like) |
| 158 | Advanta | ACSQ5-2 | 524 | BA.1 | Omicron (BA.1-like) |
| 159 | Advanta | ACSQ5-25 | 504 | BA.1 | Omicron (BA.1-like) |
| 160 | Advanta | ACSQ5-21 | 502 | BA.1 | Omicron (BA.1-like) |
| 161 | Advanta | ACSQ5-3 | 492 | BA.1 | Omicron (BA.1-like) |
| 162 | Advanta | ACSQ4-17 | 490 | B.1.1.529 | Omicron (B.1.1.529-like) |
| 163 | Advanta | ACSQ5-28 | 489 | BA.1 | Omicron (BA.1-like) |
| 164 | Advanta | ACSQ5-20 | 476 | BA.1 | Omicron (BA.1-like) |
| 165 | Advanta | ACSQ5-18 | 474 | BA.1 | Omicron (BA.1-like) |
| 166 | Advanta | ACSQ5-15 | 466 | BA.1 | Omicron (BA.1-like) |
| 167 | Advanta | ACSQ5-11 | 451 | BA.1 | Omicron (BA.1-like) |
| 168 | Advanta | ACSQ2-15 | 446 | AY.3 | Delta (B.1.617.2-like) |
| 169 | Advanta | ACSQ3-9 | 444 | AY.25 | Delta (B.1.617.2-like) |
| 170 | Advanta | ACSQ3-11 | 435 | AY.25 | Delta (B.1.617.2-like) |
| 171 | Advanta | ACSQ3-17 | 427 | AY.3 | Delta (B.1.617.2-like) |
| 172 | Advanta | ACSQ5-4 | 425 | AY.103 | Delta (B.1.617.2-like) |
| 173 | Advanta | ACSQ5-9 | 420 | BA.1 | Omicron (BA.1-like) |
| 174 | Advanta | ACSQ3-16 | 418 | AY.3 | Delta (B.1.617.2-like) |
| 175 | Advanta | ACSQ3-30 | 410 | AY.39.1 | Delta (B.1.617.2-like) |
| 176 | Advanta | ACSQ5-19 | 401 | BA.1 | Omicron (BA.1-like) |
| 177 | Advanta | ACSQ3-14 | 396 | AY.3 | Delta (B.1.617.2-like) |
| 178 | Advanta | ACSQ5-10 | 392 | BA.1 | Omicron (BA.1-like) |
| 179 | Advanta | ACSQ3-15 | 385 | AY.3 | Delta (B.1.617.2-like) |
| 180 | Advanta | ACSQ3-28 | 383 | AY.39.1 | Delta (B.1.617.2-like) |
| 181 | Advanta | ACSQ5-17 | 375 | BA.1 | Probable Omicron (BA.1-like) |
| 182 | Advanta | ACSQ3-29 | 371 | AY.39.1 | Delta (B.1.617.2-like) |
| 183 | Advanta | ACSQ2-3 | 371 | AY.25 | Delta (B.1.617.2-like) |
| 184 | Advanta | ACSQ2-6 | 371 | AY.3 | Delta (B.1.617.2-like) |
| 185 | Advanta | ACSQ2-1 | 364 | AY.25 | Delta (B.1.617.2-like) |
| 186 | Advanta | ACSQ2-9 | 349 | AY.3 | Delta (B.1.617.2-like) |
| 187 | Advanta | ACSQ2-11 | 345 | AY.3 | Delta (B.1.617.2-like) |
| 188 | Advanta | ACSQ2-8 | 338 | AY.3 | Delta (B.1.617.2-like) |
| 189 | Advanta | ACSQ3-18 | 336 | AY.3 | Delta (B.1.617.2-like) |
| 190 | Advanta | ACSQ2-22 | 330 | AY.39.1 | Delta (B.1.617.2-like) |
| 191 | Advanta | ACSQ2-10 | 316 | AY.3 | Delta (B.1.617.2-like) |
| 192 | Advanta | ACSQ2-20 | 310 | AY.39.1 | Delta (B.1.617.2-like) |
| 193 | Advanta | ACSQ2-13 | 309 | AY.3 | Delta (B.1.617.2-like) |
| 194 | Advanta | ACSQ2-7 | 305 | AY.3 | Delta (B.1.617.2-like) |
| 195 | Advanta | ACSQ3-10 | 303 | AY.25 | Delta (B.1.617.2-like) |
| 196 | Advanta | ACSQ2-12 | 302 | AY.3 | Delta (B.1.617.2-like) |
| 197 | Advanta | ACSQ5-8 | 301 | AY.103 | Delta (B.1.617.2-like) |
| 198 | Advanta | ACSQ3-25 | 298 | AY.103 | Delta (B.1.617.2-like) |
| 199 | Advanta | ACSQ2-21 | 295 | AY.39.1 | Delta (B.1.617.2-like) |
| 200 | Advanta | ACSQ4-21 | 290 | AY.111 | Delta (B.1.617.2-like) |
| 201 | Advanta | ACSQ2-2 | 289 | AY.25 | Delta (B.1.617.2-like) |
| 202 | Advanta | ACSQ3-26 | 275 | AY.103 | Delta (B.1.617.2-like) |
| 203 | Advanta | ACSQ2-17 | 265 | AY.103 | Delta (B.1.617.2-like) |
| 204 | Advanta | ACSQ2-19 | 263 | AY.103 | Delta (B.1.617.2-like) |
| 205 | Advanta | ACSQ3-27 | 247 | AY.103 | Delta (B.1.617.2-like) |
| 206 | Advanta | ACSQ3-24 | 242 | AY.3 | Delta (B.1.617.2-like) |
| 207 | Advanta | ACSQ3-22 | 241 | AY.3 | Delta (B.1.617.2-like) |
| 208 | Advanta | ACSQ1-12 | 239 | B.1 | Non- VOC |
| 209 | Advanta | ACSQ2-18 | 237 | AY.103 | Delta (B.1.617.2-like) |
| 210 | Advanta | ACSQ3-23 | 227 | AY.3 | Delta (B.1.617.2-like) |
| 211 | Advanta | ACSQ2-14 | 226 | AY.3 | Delta (B.1.617.2-like) |
| 212 | Advanta | ACSQ2-16 | 224 | AY.3 | Delta (B.1.617.2-like) |
| 213 | Advanta | ACSQ1-4 | 222 | AY.3 | Delta |
| 214 | Advanta | ACSQ1-5 | 218 | AY.3 | Delta |
| 215 | Advanta | ACSQ2-15 | 165 | AY.3 | Delta (B.1.617.2-like) |
| 216 | Advanta | ACSQ3-19 | 149 | AY.82 | Delta (B.1.617.2-like) |
| 217 | Advanta | ACSQ3-20 | 132 | None |  |
| 218 | Advanta | ACSQ1-3 | 128 | AY.3 | Delta |
| 219 | Advanta | ACSQ2-11 | 120 | AY.3 | Delta (B.1.617.2-like) |
| 220 | Advanta | ACSQ1-2 | 114 | AY.25 | Delta (B.1.617.2-like) |
| 221 | Advanta | ACSQ2-13 | 107 | AY.3 | Delta (B.1.617.2-like) |
| 222 | Advanta | ACSQ1-7 | 106 | AY.3 | Delta |
| 223 | Advanta | ACSQ3-21 | 85 | AY.3 | Delta (B.1.617.2-like) |
| 224 | Advanta | ACSQ1-6 | 58 | B.1.617.2 | Delta |
| 225 | Advanta | ACSQ1-10 | 56 | B.1.574 | Non- VOC |
| 226 | Advanta | ACSQ9 | 1597 | BA.4.6 | Omicron (BA.4-like) |
| 227 | Advanta | ACSQ9 | 2000 | BA.5.2 | Omicron (BA.5-like) |
| 228 | Advanta | ACSQ9 | 1718 | BA.5.1.1 | Omicron (BA.5-like) |
| 229 | Advanta | ACSQ9 | 2300 | BA.5.2.1 | Omicron (BA.5-like) |
| 230 | Advanta | ACSQ9 | 3708 | BA.5.1 | Omicron (BA.5-like) |
| 231 | Advanta | ACSQ9 | 2483 | BA.5.1 | Probable Omicron (BA.5-like) |
| 232 | Advanta | ACSQ9 | 2414 | BE.1.1 | Omicron (BA.5-like) |
| 233 | Advanta | ACSQ9 | 1781 | BA.5.2 | Omicron (BA.5-like) |
| 234 | Advanta | ACSQ9 | 3484 | BE.1.1 | Omicron (BA.5-like) |
| 235 | Advanta | ACSQ9 | 5257 | BA.5.2 | Probable Omicron (BA.5-like) |
| 236 | Advanta | ACSQ9 | 6032 | BA.4.6 | Omicron (BA.4-like) |
| 237 | Advanta | ACSQ9 | 5435 | BA.4.6 | Omicron (BA.4-like) |
| 238 | Advanta | ACSQ9 | 3008 | BA.4.6 | Omicron (BA.4-like) |
| 239 | Advanta | ACSQ9 | 2788 | BA.5.2 | Probable Omicron (BA.5-like) |
| 240 | Advanta | ACSQ9 | 5076 | BA.5.2 | Omicron (BA.5-like) |
| 241 | Advanta | ACSQ9 | 6913 | BA.5.5 | Omicron (BA.5-like) |
| 242 | Advanta | ACSQ9 | 2373 | BA.5.6 | Omicron (BA.5-like) |
| 243 | Advanta | ACSQ9 | 3744 | BA.5.1.6 | Omicron (BA.5-like) |
| 244 | Advanta | ACSQ9 | 3032 | BA.5.2 | Probable Omicron (BA.5-like) |
| 245 | Advanta | ACSQ9 | 3245 | BA.5.2.1 | Omicron (BA.5-like) |
| 246 | Advanta | ACSQ9 | 1066 | BA.5.5 | Omicron (BA.5-like) |
| 247 | Advanta | ACSQ9 | 3355 | BA.2.3 | Omicron (BA.2-like) |
| 248 | Advanta | ACSQ9 | 4023 | BA.5.2.1 | Probable Omicron (BA.5-like) |
